# RhoA effectors ROCK and SLK/LOK oppositely regulate VE-cadherin mobility at endothelial junctions

**DOI:** 10.64898/2026.10.28.762575

**Authors:** Tyler Nelson, Qiyao Lin, Zoya Quddoos, Henry Gatica-Gutierrez, Sofía Vargas-Hernández, Arnold Hayer, Anna-Karin Gustavsson

## Abstract

Cellular contractile forces regulate VE-cadherin dynamics, adhesion, and endothelial junction integrity. RhoA controls these forces through two parallel effector kinases: ROCK, which activates non-muscle myosin II motors, and SLK/LOK, which promote actin-membrane attachment through phosphorylation of ezrin, radixin, moesin proteins (ERM). How these two RhoA effector pathways differentially affect VE-cadherin dynamics at junctions is not known. Using single-molecule tracking of Halo-tagged VE-cadherin, we obtained a detailed characterization of VE-cadherin diffusive behavior inside and outside junctional regions in cultured human umbilical vein endothelial cells (HUVEC). VE-cadherin was substantially less mobile and more confined within junctions than in non-junctional membrane. Inhibiting ROCK with Y-27632 increased VE-cadherin mobility both inside and outside junctions, consistent with RhoA/ROCK-driven contractility restraining cadherin diffusion globally. Unexpectedly, inhibiting SLK/LOK produced the opposite effect, selectively reducing VE-cadherin mobility within junctions with no comparable change outside them. This effect was accompanied by an increase in reticular junctions, suggesting that SLK/LOK inhibition alters VE-cadherin mobility indirectly, through a change in junctional architecture, rather than through a direct effect of actin-membrane coupling on VE-cadherin itself. These findings identify SLK/LOK as a regulator of VE-cadherin dynamics acting in parallel to ROCK downstream of RhoA and point to junctional architecture as a previously unappreciated determinant of VE-cadherin mobility at endothelial cell-cell contacts.

**STATEMENT OF SIGNIFICANCE:** Endothelial adherens junctions rely on the highly regulated dynamics of VE-cadherin, yet how the nanoscale mobility of these calcium-dependent adhesion proteins is affected by actin cytoskeletal regulation remains unclear. Using fluorescence single-molecule tracking, we investigated the effects of two RhoA effector kinase pathways – ROCK (myosin-based contraction) and SLK/LOK (actin-membrane attachment) – on VE-cadherin diffusion. Surprisingly, these pathways act oppositely on VE-cadherin mobility at junctions. This suggests that junctional architecture, rather than actin-membrane coupling alone, acts as an additional determinant of VE-cadherin dynamics. Our approach sheds new light on the complex interplay between cytoskeleton regulation and endothelial junction behavior, which has potential implications in the study and treatment of important vascular and immune-related diseases.

## INTRODUCTION

Vascular endothelial cells maintain blood vessel integrity by forming a monolayer of tightly packed cells using cell-cell adhesive structures [1,2]. One such structure, the adherens junction, is vital for the dynamic remodeling of endothelial cells during blood vessel formation (angiogenesis) [3,4], as well as the selective permeability of the vessel lumen monolayer to leukocytes, which must be able to pass between the cells of the monolayer during immune inflammatory response [5–8]. Adherens junctions are formed by cadherin-family transmembrane proteins which homodimerize with cadherins from adjacent cells in a calcium-dependent (Ca^2+^) manner [3,9–11]. The cadherins associate with supportive proteins called catenins, which bind the cadherins to actin filaments, stabilizing the adherens junction and maintaining cell-cell adhesion [1,12,13].

VE-cadherin (also known as cadherin-5 or CD144) is an endothelial-specific transmembrane cell-cell adhesion receptor [14,15]. VE-cadherin serves important mechanical [16], developmental [14,17], and regulatory [3,18,19] roles in vascular endothelial cells, with implications in a variety of diseases and disorders, including inflammatory bowel disease [20], cerebral cavernous malformation [8], and cancer [8], with potential involvement in many others [21–23]. The structure of VE-cadherin consists of an extracellular part with five repeated cadherin domains (EC1-EC5), a transmembrane domain, and a cytoplasmic domain, through which it recruits intracellular adaptors. These include plakoglobin (also known as ɣ-catenin) [24,25], p-120 catenin [26–28], and β-catenin [28,29], and indirectly α-catenin via plakoglobin or β-catenin [30,31]. VE-cadherins cluster laterally via *cis*-oligomerization of the EC1 and EC2 domains, with EC3 and EC4 also being implicated in clustering [18,32]. This lateral clustering is enhanced by p-120 catenin binding and is thought to support junctional function by increasing localization of VE-cadherin at the plasma membrane and thus the avidity of cell-cell binding interactions [33]. Intercellular binding occurs between VE-cadherin of neighboring cells via strand-swap *trans*-dimerization of opposing EC1 domains [34].

VE-cadherin and many of the other parts of the adherens junction are closely associated with the actin cytoskeleton (Figure 1A,B), which has been known to affect the diffusion of membrane-bound proteins in a variety of ways [35–39]. Importantly, actin is also intrinsically linked to endothelial cell contractility. The contractility of endothelial cells is a major determinant of vascular function by regulating barrier permeability [40] and vascular remodeling [41], and is regulated in large part by the GTPase RhoA (and subfamily members RhoB, RhoC) via non-muscle myosin-II (NM2) and the ezrin, radixin, and moesin proteins (ERM) [42–45]. NM2 drives contractility of actin stress fibers upon activation by Rho-associated kinase (ROCK) [46–48]. ERM, upon activation by the RhoA effectors Ste20-like kinase (SLK) and lymphocyte-oriented kinase (LOK), link cortical actin to the plasma membrane [49–53]. ROCK and SLK/LOK thus represent two parallel RhoA effector arms that organize cellular contractility by mechanistically distinct means – direct NM2-based motor activity versus actin-membrane attachment. Studying how the perturbation of these kinases, ROCK and SLK/LOK, affect VE-cadherin dynamics can inform how cytoskeletal forces and configurations modulate VE-cadherin and adherens junction behavior.

**Figure 1:**
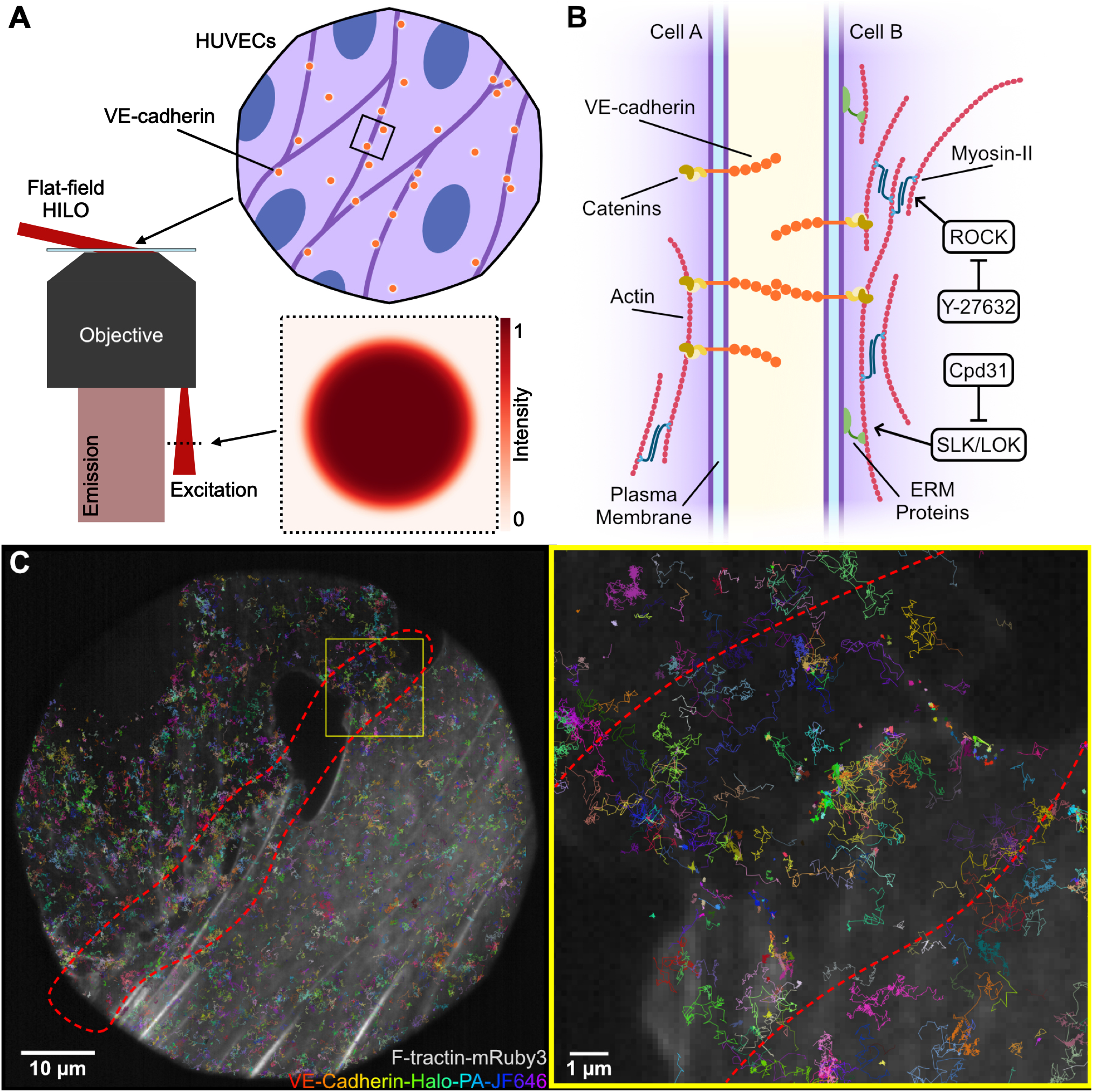
The dynamics of single VE-cadherin-HaloTag molecules were tracked in HT-HUVEC with and without perturbations to RhoA effector pathways. (A) Schematic of the flat-field HILO illumination configuration used for single-particle tracking of VE-cadherin-HaloTag in HT-HUVEC. (B) Schematic of an endothelial adherens junction, highlighting the relationship between VE-cadherin, the actin cortex, and the plasma membrane, as well as the RhoA effector pathways ROCK and SLK/LOK and inhibitors Y-27632 and Cpd31, respectively. (C) Fluorescence image of HT-HUVEC showing the actin cytoskeleton (Ftractin-mRuby3) with VE-cadherin-HaloTag trajectories overlaid. Right: Magnified inset of boxed area in left image showing details of part of a cell-cell junction and the variety of trajectories present. Colors represent distinct VE-cadherin-HaloTag trajectories. Red dashed line represents the manual selection of the area in proximity to a cell-cell junction, which was used to categorize trajectories during analysis.

Despite VE-cadherin’s crucial functions, how its dynamics are shaped by these two RhoA effector pathways remains unknown. To address this gap, we investigated how targeted disruption of actin-regulating pathways affects VE-cadherin behavior. Specifically, we examined the effects of a ROCK inhibitor, Y-27632, and an SLK/LOK inhibitor, Cpd31, on the dynamics of VE-cadherin-HaloTag in hTERT immortalized human umbilical vein vascular endothelial cells (HT-HUVEC) (Figure 1A,B). We performed single-particle tracking (SPT) on individual VE-cadherin molecules diffusing in the plasma membrane, with nanometric spatial and millisecond temporal resolution (Figure 1C). The VE-cadherin was categorized based on location within the cell, and multiple characteristics were compared before and after treatment. We first established that VE-cadherin mobility is reduced within junctions relative to non-junctional membrane, consistent with confinement imposed by trans-dimerization and cortical actin coupling. We showed that inhibition of ROCK, which is responsible for actin stabilization and NM2 contraction, allowed VE-cadherin to diffuse more freely and explore larger areas. We furthermore demonstrated that, unexpectedly, inhibition of SLK/LOK, activators of ERM and actin membrane attachment, led to more restricted diffusion of VE-cadherin near cell-cell junctions, while not significantly affecting VE-cadherin localized far from junctions. These findings highlight a previously unrecognized complexity in the relationship between actin organization and VE-cadherin diffusion, revealing how actin-regulatory pathways modify VE-cadherin mobility which may ultimately modulate adherens junction behavior.

## MATERIALS AND METHODS

### Cell culture and generation of cells stably expressing reporter constructs

hTERT-immortalized HUVEC (HT-HUVEC), from mixed (male, female) donors [54], were cultured in endothelial cell growth medium (PromoCell, C-22216 and C-39216, or C-22011), supplemented with hygromycin (50 μg/mL; Invivogen, ant-hg-1). HT-HUVEC stably expressing reporter constructs were established via lentiviral transduction followed by selection with the appropriate antibiotic puromycin (0.5 μg/mL; Invivogen, ant-pr-1) and G418 (0.5 mg/mL; Invivogen, ant-gn-2). Generation of cells co-expressing VE-cadherin-HaloTag and Ftractin-mRuby3 was achieved by sequential lentiviral transductions and corresponding antibiotic selections. Stably transduced cells were maintained in the presence of puromycin and G418. The cell suspension was split as needed into new flasks or 8-well chambered coverslips (Ibidi, 80827). Flasks and chambered coverslips were all coated with type I bovine collagen (Advanced Biomatrix, #5005) by incubating at 37°C with 1% vol/vol collagen in PBS for 30 min, followed by a wash with PBS. Cells were seeded in chambered coverslips 1 day prior to imaging.

### DNA constructs

The pLV-CAG-CDH5-HaloTag-IRES-puro lentiviral vector was generated by replacing the mCitrine sequence in pLV-CDH5-mCitrine-IRES-Puro (Addgene, #85143 [54]) with the HaloTag sequence. The HaloTag sequence was PCR-amplified from pUC-HaloTag-Kan (kindly provided by Dr. Rodrigo Reyes) and inserted into the NotI/Agel-digested pLV-CDH5-mCitrine-IRES-Puro using Gibson assembly.

pLV-Ftractin-mRuby3-IRES-Neo was created by PCR-amplifying Ftractin-mRuby3 using pLV-Ftractin-mRuby3-p2A-mTurquoise-MLC-IRES-Blast as template (Addgene, #85146 [54]) and by inserting the product into BamHI/NotI-digested pLV-EF1a-MCS-IRES-Neo (Addgene, #85139 [54]) using Gibson assembly.

### Production of lentivirus

Lentiviral particles were produced in HEK293FT cells (not authenticated). HEK293FT cells were cultured in DMEM (Thermo Fisher Scientific, 11965092) supplemented with 10% fetal bovine serum (FBS; Corning, 35-077-CV) and 1% GlutaMAX (Thermo Fisher Scientific, 35050061). To package the virus, cells were co-transfected with lentiviral transfer plasmids and the third-generation packaging plasmids (pMDLg/pRRE, pRSV-REV, and pCMV-VSVG) using Lipofectamine 2000, as described in [45]. Viral supernatants were harvested at 48 h post-transfection and filtered through a 0.22 μm filter (VWR CA28143-310).

### VE-cadherin labeling and imaging buffer

A HEPES-buffered, CO2-independent medium for live HT-HUVEC imaging with low background fluorescence, herein referred to as LIS (live-cell imaging solution), was used for all imaging experiments (adapted from [54]). First, extracellular buffer was prepared by combining 5 mM KCl (Sigma-Aldrich, P3911), 125 mM NaCl (Sigma-Aldrich, S9625), 20 mM HEPES (Sigma-Aldrich, 83264), 1.5 mM MgCl2 (Fisher Scientific, BP214-500), 1.5 mM CaCl2 (Thermo Scientific, J62905.AP), and 10 mM D-glucose (Sigma-Aldrich, G5767), then adjusting pH to 7.4 using HCl or KOH. Prior to imaging, fetal bovine serum (FBS, Gibco, 16-000-044) and recombinant human basic fibroblast growth factor (bFGF, R&D Systems, 233-FB) were added such that the resulting solution contained 1% FBS and 5 ng/mL bFGF.

For SPT, photoactivatable PA-JF646-HaloTag dye (Janelia Research Campus, [55]) was diluted from a 50 µM aliquot to 1 nM in LIS at 37°C. HT-HUVEC were first washed once with 37°C Mg- and Ca-free PBS, then incubated in the dye solution for 30 min at 37°C. The dye solution was then aspirated, and the cells were washed for 15-30 s three times with 37°C PBS. Wells were refilled with LIS, and the cells were placed back in an incubator at 37°C for 15 min prior to imaging. Protocol adapted from [56].

### Drug treatments

For all live-cell drug treatment experiments, the same FOVs were imaged both before and after treatment. Cell media was then fully aspirated, and the drug compound was added to the well in LIS. For ROCK inhibition experiments, cells were treated with 20 µM Y-27632 (MedChemExpress, HY- 10583R) for 1 h, then imaged immediately after. For SLK/LOK inhibition experiments, cells were treated with 5 µM Cpd31 (MedChemExpress, HY-132868) for 15 min, then imaged. For drug treatment vehicle control experiments, cells were incubated with 20 µM DMSO (Sigma-Aldrich, D2438-5X10ML) for 1 h, then imaged. For fixed-cell imaging, cells were control-treated or treated with 20 µM Y-27632 or 5 µM Cpd31 for 2 h prior to fixation.

### Immunolabeling and fixed-cell imaging

Cells were plated at 15,000 cells/well in collagen-coated glass-bottom 96-well plates (Cellvis P96-1.5H-N) and cultured overnight. Following control or drug treatments, cells were fixed using 4% formaldehyde (Fisher Scientific, 50-980-487) in PBS for 15 min at room temperature (RT), then rinsed with PBS. Permeabilization and blocking was done using ASBB (10% FBS (Fisher Scientific, 35077CV), 1% bovine serum albumin (BSA; BioShop Canada, ALB001.250), 0.1% Triton X-100 (MilliporeSigma, T9284), and 0.01% NaN3 (BioShop Canada, SAZ001)) for 30 min, followed by incubation with anti-VE-cadherin primary antibodies (BD Biosciences, 610252) diluted 1:400 in ASBB overnight at 4°C. Following three washes with PBS, 5 min each, cells were incubated with Alexa Fluor 647-conjugated secondary antibodies (Fisher Scientific, A21235), diluted 1:1,000 in ASBB, and phalloidin-Alexa Fluor 488 (New England Biolabs, 8878S) diluted 1:200, for 1 h at RT, followed by three washes with PBS, 5 min each. Cells were imaged overlaid with PBS using an automated epifluorescence microscopy system built around a Nikon TI-2E stand, equipped with a Lumencor SpectraX LED light source, a FLI HS-1025 emission filter wheel, a Hamamatsu Orca Fusion-BT camera, and a 40x 1.3 NA CFI Plan Fluor objective.

### Optical setup for live-cell imaging

A custom-built microscope for live-cell imaging with homogeneous epi-/TIRF illumination, which ensures uniform photophysics across the field of view, was used to acquire SPT data [57]. After cells were labeled, immersion oil (Zeiss, 444970-9010-000) was placed on the objective, then an 8-well chambered coverslip containing HT-HUVEC coexpressing the F-actin marker Ftractin-mRuby3 and VE-cadherin-HaloTag was placed in the stage-top incubator (Okolab, OKO-H301PI736ZR1/2S). Focus was found using low-intensity (∼73 mW/cm^2^) 560 nm (MPB Communications, 2RU-VFL-P-1000-560-FC) epi-fluorescence, then the microscope’s TruFocus Z-drift compensation system (Olympus, IX3SP-ZDC2-830) was activated, locking the focus. The sample was then left to settle for 15 min to reduce sample drift while imaging. The flat-field epi-/TIRF stage was set to a point between epi- and TIRF to achieve flat-field HILO illumination [58] for improved range and contrast. 560 nm laser illumination at ∼73 mW/cm^2^ was used for actin imaging. Diffraction-limited actin fluorescence images were acquired just before and after SPT acquisition for each field of view (FOV) and consisted of one hundred 25 ms frames, which were then averaged together. For VE-cadherin tracking acquisition, the 405 nm laser (Coherent, 1284371) was set to ∼11 mW/cm^2^ using a tunable optical density filter. This laser was used to activate the photoactivatable dye. The 642 nm laser (MPB Communications, 2RU-VFL-P-1000-642-FC), used to excite the photoactivatable dye, was set for 80 W/cm^2^ intensity. Both lasers were used in unison throughout the 7500-frame acquisition for each FOV. Laser intensities were chosen to balance the number of activated fluorophores with the number of photobleached fluorophores in each frame, while achieving optimal contrast at 25 ms exposure times (40 frames per second) with 2x2 pixel binning, standard readout speed settings, and a 100X, 1.5 NA objective (Olympus, UPLAPO100XOHR).

### Single-particle tracking analysis

Images of VE-cadherin-HaloTag molecules were analyzed in TrackMate [59,60] to extract their dynamics parameters. In TrackMate, a Laplacian of Gaussian filter with σ = 0.5 µm and quadratic maxima fitting for sub-pixel localization was used to identify the positions of single fluorophores. To choose a threshold for localization, the quality histogram was assessed. The quality measurement represents the local maxima of the filtered image, where brighter spot detections have correspondingly higher quality. There were typically four distinct populations of potential localizations in this histogram. The lowest quality population corresponded to areas of the camera sensor that were outside the bounds of the circular channel and were thus not illuminated. The second lowest quality population corresponded to parts of the channel where no cells were present. The third population consisted of out-of-focus fluorophores. The fourth, highest quality, population of potential localizations corresponded to in-focus fluorophores. The quality threshold was set to capture primarily only this population. LAP (Linear Assignment Problem) tracker with maximum linking distance of 0.4 µm was used for trajectory connection, with maximum segment gap closing distance of 0.4 µm and maximum frame gap of 10. Trajectories were filtered such that each had a minimum of 10 localizations. Further filtering (*vide infra*) increased this minimum to 40.

### Junction discrimination and selection

A custom python script was written which allowed the blinded selection of trajectories in close proximity to cell-cell junctions based on visual inspection of diffraction-limited Ftractin-mRuby3 images. The Ftractin-mRuby3 image was displayed without the corresponding trajectories, and cell-cell junctions were selected by encircling them using a mouse. Areas with obvious contact between cells were selected and each selection also included a buffer of 2-3 µm to account for any potential cell movement or drift that may have occurred during the course of the acquisition.

### Mean square displacement and scan area calculation

The discrete-time mean square displacement of a single particle was calculated according to Eq. (1) [61]:

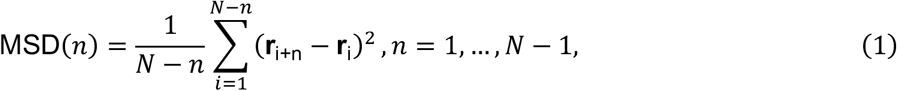

Where **r**_i+n_ is the *d*-dimensional (in this case, 2-dimensional) position of the particle at frame *i* + *n*, **r**_i_ is the reference position of the particle at frame *i*, *n* is the interval, in frames, between localizations (also known as time lag), and *N* is the total number of frames in a trajectory. Because of the large number of trajectories in each acquisition (tens of thousands), and the potential for many of those trajectories to be very long (thousands of frames), using the above-described method for calculating MSD could be computationally demanding ( α *O*(N^2^) ). To facilitate MSD calculation, the Fast Correlation Algorithm (FCA) [62] was implemented, which reduced the computational complexity to be α*O*(*N*log_2_(*N*)2.

MSDs were fitted with the expression in Eq. (2) to extract the effective diffusion coefficient *D*^∗^ , anomalous diffusion exponent α, and static error σ [63–67]:

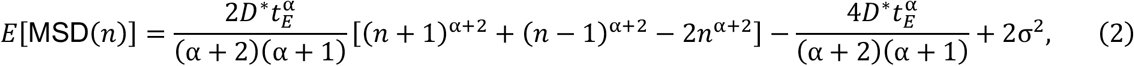

where *t_E_* is the camera exposure time and *n* is the lag time (in number of frames). Trajectories shorter than 40 frames (1 s) were discarded to ensure robust fitting statistics. Additionally, fitted parameters were constrained to prevent parameter degeneracy and unphysical results: *D*^∗^ ∈ (0, 2] µ*m*^2^/s**^α^**, α ∈ [0.0025, 1.9975], and σ ∈ [0, 1] μm.

While the MSD allows for the estimation of effective diffusion coefficient and classification of diffusion type, it cannot necessarily distinguish between two particles with similar diffusion coefficients, where one is confined to a particular area while the other is not. Calculating the scan area of a trajectory can give insight into changes in local environment experienced by VE-cadherin. For each trajectory, the convex hull area was calculated [68,69] to determine the area scanned by each tracked particle over its total length.

### Statistical Analysis

Each experiment was performed in triplicate across different days. 32 FOVs were captured in each experiment – 16 before and 16 after treatment – for a total of 48 datasets for each condition and location combination (e.g., *n* = 48 post-ROCK inhibitor non-junction datasets). Each FOV consisted of 7500 frames and resulted in a median of 826 trajectories. Trajectories ranged from 40 to 7500 frames, with a median length of 119 frames (2.975 s). Because the data are hierarchical and consist of many replicates, a linear mixed effects model [70] was used to test for statistical significance of the fixed effect (drug treatment), with variable slope and intercepts to account for random variation in cell samples and their response to treatment. P-values were calculated with Satterthwaite’s method using the “lmerTest” package in R [71].

## RESULTS

### ROCK and SLK/LOK inhibition differentially alter VE-cadherin distribution at cell-cell junctions

To establish context for how RhoA effector kinase inhibition affects VE-cadherin organization prior to investigating nanoscale molecular dynamics, we first visualized the organization of F-actin and VE-cadherin in HT-HUVEC using epifluorescence microscopy following fixation and immunofluorescence staining (Figure 2, Supplemental Figure S1-S3). We used HT-HUVEC that stably expressed the F-actin marker Ftractin-mRuby3 and VE-cadherin-HaloTag. In control-treated cells (Figure 2A), VE-cadherin exhibited typical behavior, with primarily well-defined, highly clustered, sometimes finger-like, localization along cell-cell contacts. In some areas, reticular junctions [72] were observed, particularly where cells seem to partially “subduct” beneath neighboring cells. Inhibition of ROCK with Y-27632 resulted in a visual reduction of actin stress fiber size and a thinning of VE-cadherin in cell-cell junctions, with VE-cadherin localization appearing less finger-like and junctions less tortuous than in control cells (Figure 2B). A similar number of reticular junctions can be observed as in the control images. Cells treated with the SLK/LOK inhibitor Cpd31 showed a marked increase in the prevalence of reticular junctions compared to control cells, especially in areas with long, contiguous, cell-cell contacts (Figure 2C). These results indicate that the two RhoA effector pathways differentially regulate junctional architecture, with ROCK inhibition altering VE-cadherin distribution along existing junctions and SLK/LOK inhibition instead promoting the formation of reticular junctions.

**Figure 2:**
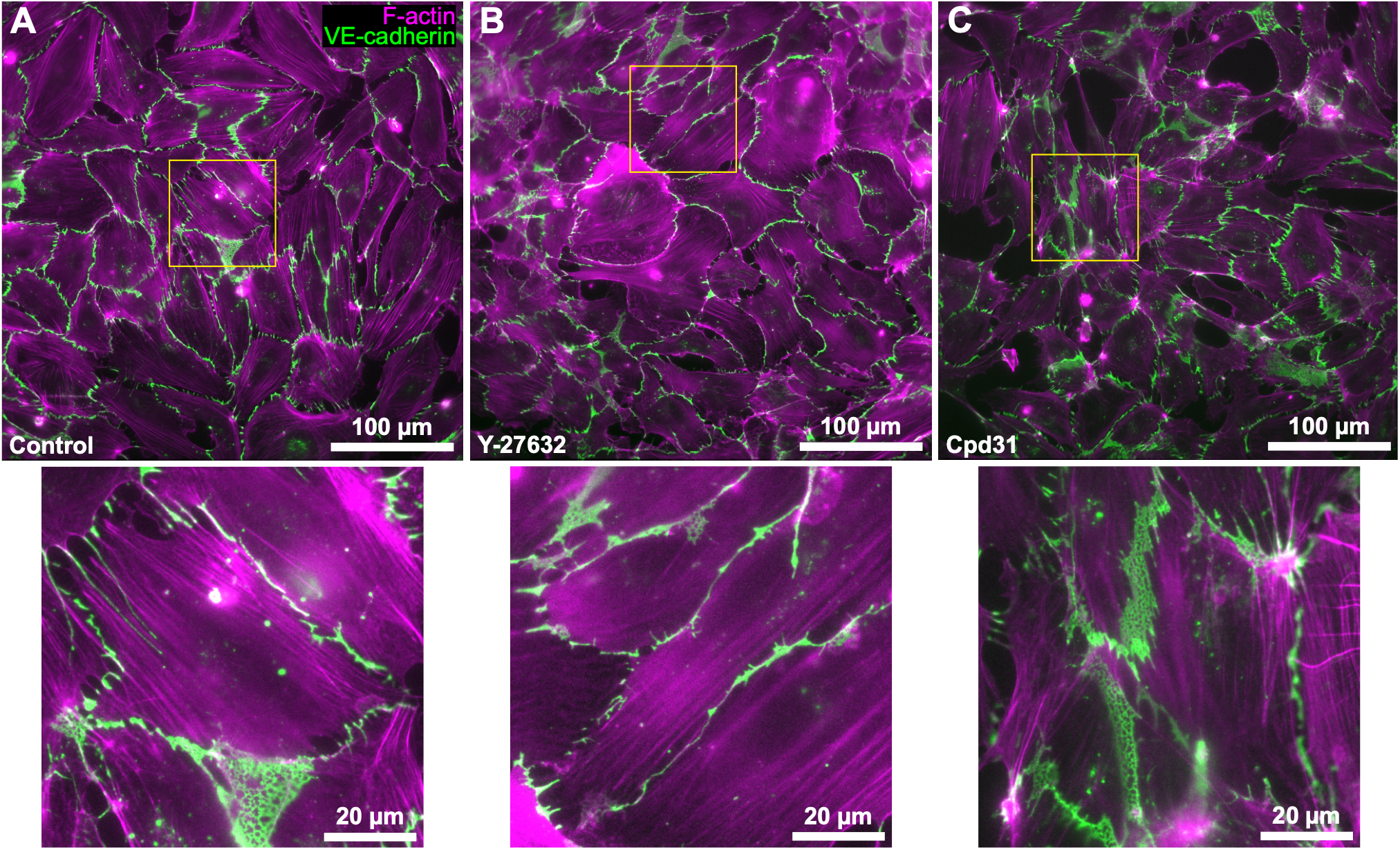
Representative epifluorescence images of HT-HUVEC co-expressing Ftractin-mRuby3 and VE-cadherin-HaloTag labeled for F-actin (phalloidin, magenta) and VE-cadherin (anti-VE-cadherin, green), all treated for 2 h prior to fixation. (A) Control-treated HT-HUVEC showing a mix of junction phenotypes, with reticular junctions seen in some areas. (B) HT-HUVEC treated with the ROCK inhibitor Y-27632 do not appear to exhibit increased reticular junction formation, and cell-cell junctions appear thinner than in untreated cells. (C) HT-HUVEC treated with the SLK/LOK inhibitor Cpd31 displayed an increased prevalence of reticular junctions between cells.

### VE-cadherin diffusion is reduced within cell-cell junctions relative to non-junctional membrane

Before assessing the effects of RhoA effector kinase inhibition, we first compared baseline VE-cadherin-HaloTag diffusion inside and outside cell-cell junctions in control (pre-treatment/DMSO-treated) HT-HUVEC that stably expressed Ftractin-mRuby3 and VE-cadherin-HaloTag. In each treatment case, cells were imaged and analyzed both before and after treatment, labeled as “pre” and “post,” respectively. VE-cadherin was significantly less mobile and more confined within junctions than in non-junctional membrane, with lower effective diffusion coefficient (0.038 μm^2^/s^α^ average in junctions vs 0.063 μm^2^/s^α^ average outside junctions) and smaller scan area (7.467 μm^2^ average in junctions vs 10.690 μm^2^ average outside junctions) at junctions compared to non-junctional regions, both before and after vehicle addition (Figure 3). This difference was consistent across conditions, indicating that junctional and non-junctional VE-cadherin represent two stable, consistently distinct diffusive populations, unaffected by vehicle addition itself. This baseline distinction establishes the framework for interpreting the differential effects of ROCK and SLK/LOK inhibition described below.

**Figure 3:**
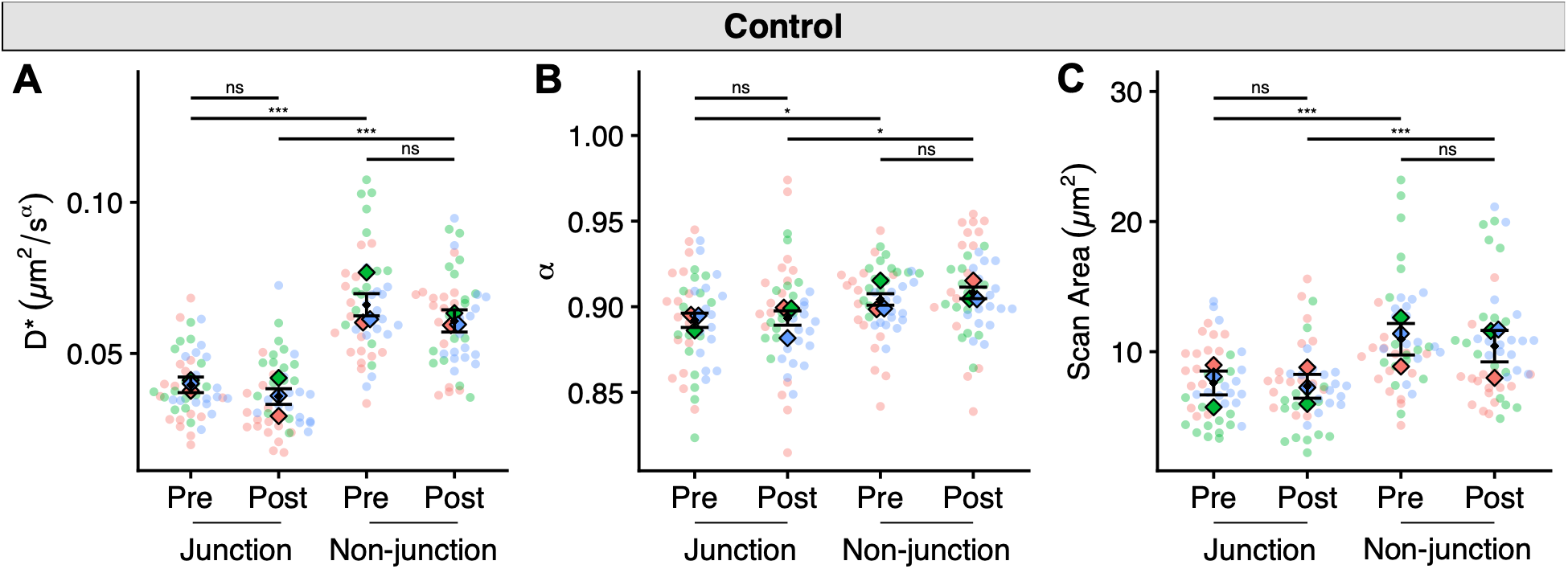
VE-cadherin-HaloTag diffusion is reduced in cell-cell junctions. Results of single-particle tracking analysis of VE-cadherin diffusion in HT-HUVEC before (Pre) and after (Post) treatment with DMSO only, grouped by the area analyzed (Junction or Non-junction). DMSO concentration matched that used for Y-27632 treatment (Fig. 4), which used a higher concentration of DMSO than the Cpd31 treatment (Fig. 5). Non-junctional VE-cadherin-HaloTag display consistent, significantly increased mobility compared to junctional VE-cadherin-HaloTag. (A) Effective diffusion coefficient, (B) anomalous exponent, and (C) scan area. For each plot, circular data points represent the median value from one field of view. Colors distinguish between different biological replicates of the same experiment. Diamond-shaped data points represent the mean value of a biological replicate. Error bars show the pooled standard error of the mean of fixed effects, centered on the estimated marginal mean of each condition. ns: p>.05, *: p<0.05, **: p<0.01, ***: p<0.001.

### ROCK inhibition increases VE-cadherin diffusion and explored area throughout the cell

ROCK acts downstream of RhoA and affects cytoskeletal organization by increasing activation of NM2, thereby increasing cytoskeletal forces. To probe the effects of decreased cytoskeletal forces on VE-cadherin diffusion, we treated HT-HUVEC with the ROCK inhibitor Y-27632. After 1 h of treatment with Y-27632, VE-cadherin-HaloTag trajectories displayed freer movement overall compared to measurements before treatment, as well as to control treatment with only DMSO (**Error! Reference source not found.**Figure 3), regardless of whether they were in the vicinity of a cell-cell junction or not (Figure 4). Near junctions, VE-cadherin-HaloTag had significantly higher effective diffusion coefficient (0.040 to 0.052 µ*m*^2^/*s^a^*, Figure 4A) and scan area (7.11 to 9.18 µ*m*^2^, Figure 4C) after treatment. In non-junctional regions, VE-cadherin displayed a similar, slightly larger, increase in effective diffusion coefficient (0.062 to 0.079 µ*m*^2^/*s^a^*, Figure 4A) and scan area (9.98 to 15.42 µ*m*^2^, Figure 4C). Notably, while the diffusion coefficient and scan area increased at and away from junctions, the difference between junctions and non-junctions remained. The two populations of VE-cadherin remained distinct, with junctional VE-cadherin showing slower diffusion. Near junctions, α did not show a significant change after treatment (Figure 4B). Away from junctions, α was significantly higher after treatment, though in absolute terms the difference was rather small (α = 0.903 vs α = 0.915, Figure 4B). Overall, ROCK inhibition increased the diffusion coefficient and area scanned by VE-cadherin throughout the cell.

**Figure 4:**
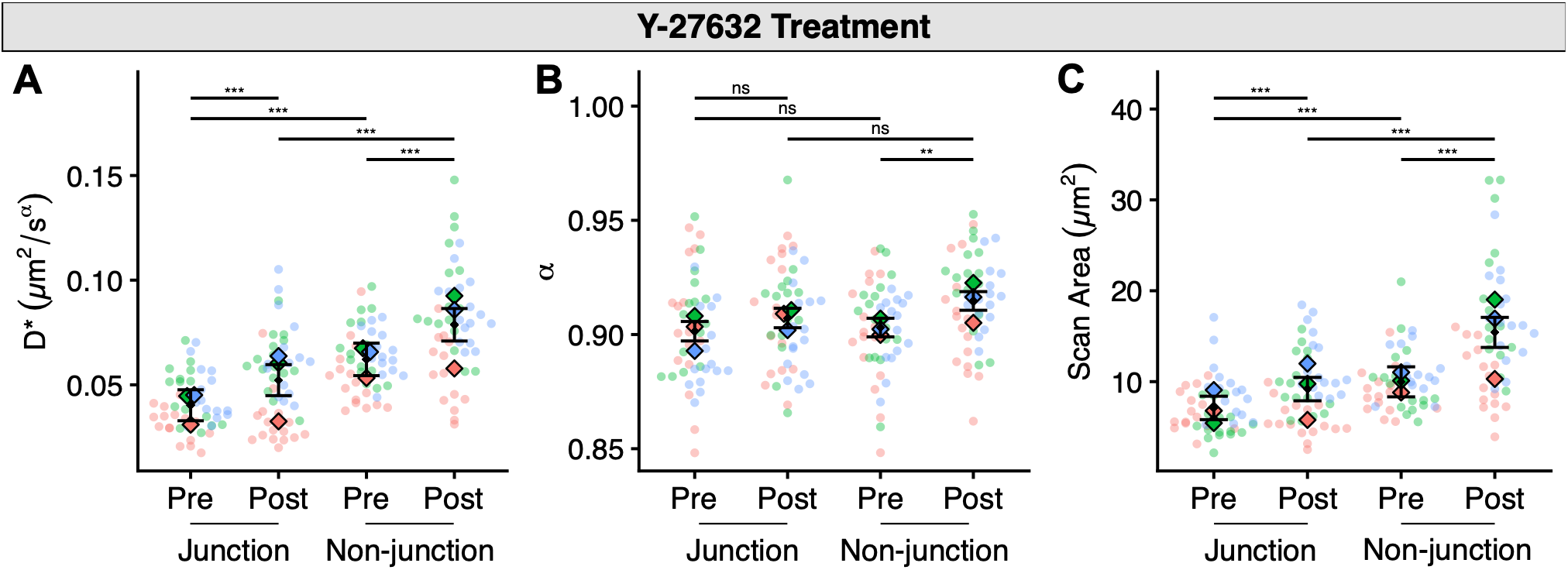
VE-cadherin-HaloTag diffusion increases across the cell following ROCK inhibition. Results of single-particle tracking analysis of VE-cadherin-HaloTag diffusion in HT-HUVEC before (Pre) and after (Post) treatment with 20 μM ROCK inhibitor Y-27632 for 1 h, grouped by the region analyzed (Junction or Non-junction). (A) Effective diffusion coefficient, (B) anomalous exponent, and (C) scan area. For each plot, circular data points represent the median value from one field of view. Colors distinguish between different biological replicates of the same experiment. Diamond-shaped data points represent the mean value of a biological replicate. Error bars show the pooled standard error of the mean of fixed effects, centered on the estimated marginal mean of each condition. ns: p>0.05, *: p<0.05, **: p<0.01, ***: p<0.001.

### SLK/LOK inhibition restricts VE-cadherin diffusion primarily in cell-cell junctions

To assess the relationship between VE-cadherin dynamics and the linkage of the actin cortex to the membrane, we treated HT-HUVEC with the SLK/LOK inhibitor Cpd31 [45,73] (Figure 5). SLK/LOK act downstream of RhoA to activate ERM [49–51]. Upon treatment, VE-cadherin-HaloTag trajectories exhibited significantly lower diffusion coefficient (0.056 to 0.040 µ*m*^2^/*s^a^*, Figure 5A) and scan areas (8.17 to 5.47 µ*m*^2^, Figure 5C) near cell-cell junctions. Trajectories outside of junctions displayed slightly lower scan area (13.39 to 10.87 µ*m*^2^ , Figure 5C) and no significant change in effective diffusion coefficient after treatment (Figure 5A). There was no significant change in α inside or outside of junctions (Figure 5B). These results were unexpected: rather than the increased mobility predicted from reduced membrane-cortex linkage, SLK/LOK inhibition instead produced more restricted VE-cadherin diffusion within junctions. A possible explanation is the formation of reticular junctions due to SLK/LOK inhibition. Initial observations indicate that reticular junctions are more prevalent in HT-HUVEC which have been treated with Cpd31 (Figure 2, Supplemental Figure S3).

**Figure 5:**
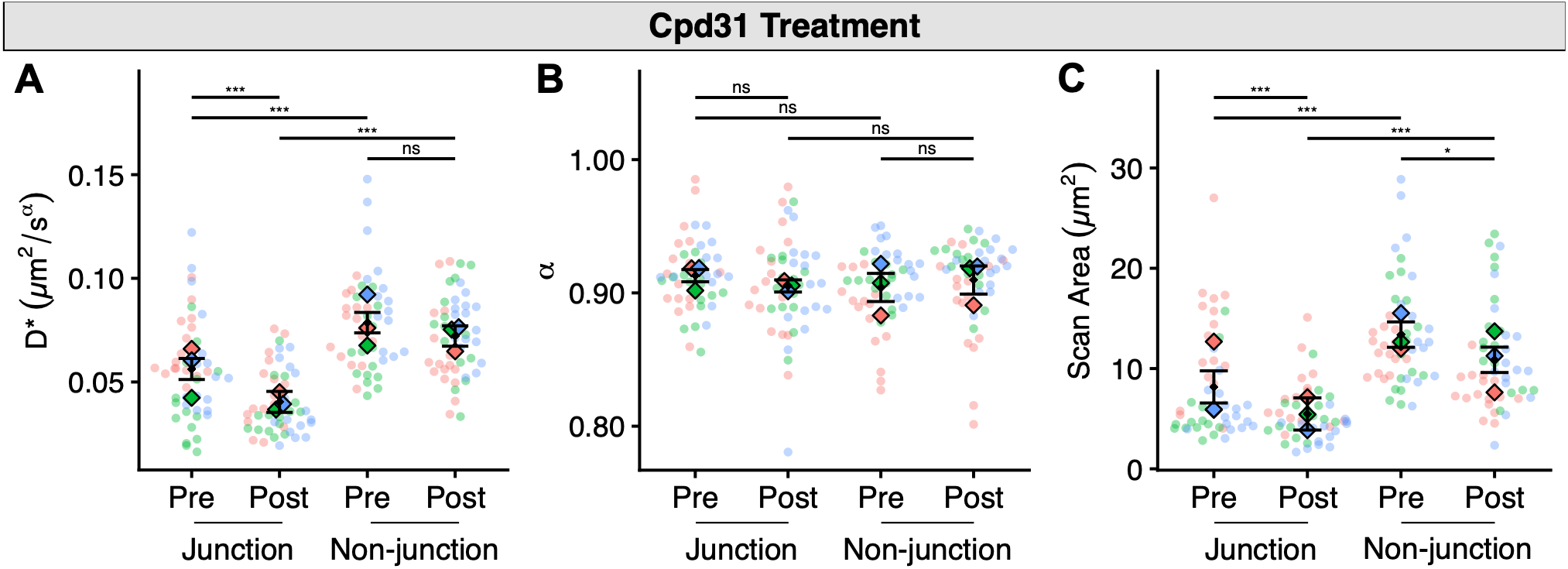
Following SLK/LOK inhibition, VE-cadherin-HaloTag dynamics are hampered primarily within cell-cell junctions. Results of single-particle tracking analysis of VE-cadherin diffusion in HT-HUVEC before (Pre) and after (Post) treatment with the SLK/LOK inhibitor Cpd31, grouped by the area analyzed (Junction or Non-junction). (A) Effective diffusion coefficient, (B) anomalous exponent, and (C) scan area. For each plot, circular data points represent the median value from one field of view. Colors distinguish between different biological replicates of the same experiment. Diamond-shaped data points represent the mean value of a biological replicate. Error bars show the pooled standard error of the mean of fixed effects, centered on the estimated marginal mean of each condition. ns: p>0.05, *: p<0.05, **: p<0.01, ***: p<0.001.

## DISCUSSION AND CONCLUSIONS

Endothelial barrier integrity and flexibility are reliant on the regulation of adherens junctions. This is a dynamic process involving the active remodeling of cortical actin, and VE-cadherin, as a transmembrane protein, should be sensitive to actin dynamics and remodeling [36]. While previous studies have established the important role of F-actin organization in VE-cadherin behavior [13,74], precisely how distinct modes of actin regulation affect the nanoscale dynamic behavior of individual VE-cadherin proteins remains underexplored [75]. We used single-particle tracking to investigate the effects of modulating cellular contractility, by acutely inhibiting upstream kinases, on VE-cadherin dynamics. These kinases are not known to directly interact with VE-cadherin – instead, they regulate the stabilization of actin and the contraction of stress fibers via NM2 in the case of ROCK, and the linkage of the actin cortex to the plasma membrane in the case of SLK/LOK. When comparing trajectories at and away from junctions, before any treatment, we observed lower effective diffusion coefficient and scan area near cell-cell junctions compared with elsewhere in the cell. We did not typically see significant differences in α , inside or outside of junctions, indicating no systemic differences in the type of diffusion. This baseline confinement of VE-cadherin at junctions likely reflects known junctional stabilization mechanisms, including trans-dimerization with VE-cadherin on adjacent cells and lateral clustering coupled to cortical actin [34,76–78]. The absence of a corresponding shift in α suggests that junctional confinement constrains the extent of VE-cadherin diffusion without altering its underlying diffusive character.

In this study, we sought to determine how the two parallel RhoA effector pathways, ROCK and SLK/LOK, differentially modulate VE-cadherin diffusion. Our results show that VE-cadherin exhibits freer diffusion after treatment with ROCK inhibitor than before treatment. ROCK inhibition reduces the activity of NM2, lowering cortical tension across the cell. This aligns with the observed increase in effective diffusion coefficient and scan area of VE-cadherin-HaloTag in ROCK-inhibited HT-HUVEC. A similar increase was measured throughout cells, not only in junctions, indicating that this loss of NM2-mediated contractility affects VE-cadherin globally, whether directly or indirectly. A small but significant increase in α was also observed outside of junctions, pointing to a potential differential effect in diffusion type when compared to trajectories inside junctions. This effect shows VE-cadherin may diffuse in a manner slightly closer to pure Brownian motion away from junctions. In all datasets, VE-cadherin diffusion was generally slower near cell-cell junctions and faster in areas further away. This difference remained after treatment with both Y-27632 and Cpd31, suggesting that both junctional engagement and F-actin linkage affect VE-cadherin dynamics.

While reducing actomyosin contractility via ROCK inhibition produced an expected increased mobility of VE-cadherin throughout the cell, weakening the physical membrane-cortex connection using the SLK/LOK inhibitor Cpd31 resulted in a seemingly paradoxical localized decrease in VE-cadherin mobility and scan area in cell-cell junctions. Outside of junctions, this effect was not observed, aside from a slight decrease in scan area. The manner in which SLK/LOK inhibition would affect VE-cadherin diffusion is not entirely clear. The most direct effect of SLK/LOK inhibition is the suppression of activation of ERM. Considering this effect, one might expect SLK/LOK inhibition to increase the effective diffusion coefficient of a membrane-bound protein like VE-cadherin. Imaging of Cpd31-treated HT-HUVEC shows an increase in reticular junctions after treatment. We hypothesize that SLK/LOK inhibition induces the formation of these convoluted, web-like structures, which restrict the movement of VE-cadherin inside junctions, while leaving VE-cadherin outside junctions largely unaffected.

A plausible mechanistic link between ERM activity and junctional architecture is the role of ERM-dependent actin-membrane attachment in controlling local protrusion-retraction dynamics at cell-cell contacts [45,79,80]. Endothelial junctions are continuously remodeled by cycles of lamellipodial protrusion and retraction, a process implicated in the consolidation of VE-cadherin into dense, junctional clusters [81–83]. ERM couple the actin cortex to the membrane and are required for the mechanical integrity of local retractions [45]; their loss could therefore be expected to dampen protrusion-retraction cycling at junctions. In this model, reduced motility at the junctional cortex following SLK/LOK inhibition would leave VE-cadherin unable to be efficiently swept into consolidated clusters, instead remaining distributed across a more disperse, reticular network - a structural change that could plausibly account for both the altered junction morphology and the reduced VE-cadherin mobility we observe. We did not directly test protrusion-retraction dynamics in this study, and this model remains to be tested, for instance by live imaging of junctional actin dynamics or lamellipodial activity under SLK/LOK inhibition.

When considered together, the findings presented in this paper reveal that ROCK and SLK/LOK, two parallel effector kinase pathways acting downstream of RhoA, differentially regulate VE-cadherin mobility at endothelial junctions. Reducing actomyosin contractility via ROCK inhibition increases VE-cadherin diffusion and scanned area globally. Conversely, disrupting membrane-cortex linking by ERM through SLK/LOK inhibition decreases the diffusion coefficient of VE-cadherin exclusively within junctions, while leaving non-junctional VE-cadherin largely unaffected. This divergence, despite a shared upstream regulator, points to junctional architecture – rather than cortical tension alone – as a previously unappreciated determinant of VE-cadherin mobility at cell-cell contacts. Our data suggest that VE-cadherin mobility, and, presumably, adherens junction functionality, reflects a regulated balance of cortical tension, NM2 contractility, and junctional architecture itself.

Future studies could use super-resolution microscopy to better characterize the structure of reticular junctions and their relationship to SLK/LOK inhibition, and live-cell imaging of junctional actin and lamellipodial dynamics could directly test whether reduced protrusion-retraction cycling underlies the reticular junction phenotype [57,84–93]. Further development of our SPT analysis pipeline could reveal more subtle effects of actin remodeling and junctional structure on VE-cadherin mobility, and how these changes affect junctional function overall.

By revealing junctional architecture as a distinct, RhoA-linked determinant of VE-cadherin mobility, this work adds a new axis to how endothelial junction function – and dysfunction – can be understood, with direct relevance to vascular permeability disorders, autoinflammatory disease, and tumor angiogenesis.

## DATA AND CODE AVAILABILITY

Custom scripts used for data analysis are available on GitHub: https://github.com/Gustavsson-Lab/ve-cadherin-analysis. Data presented in this study are available from the corresponding authors upon request.

## ACKNOWLEDGMENTS

This work was supported by the National Institute of General Medical Sciences of the National Institutes of Health grant R35GM155365 and startup funds from the Cancer Prevention and Research Institute of Texas grant RR200025 to A.-K.G., as well as by a Canadian Institutes of Health Research (CIHR) Project Grant (PJT-165932) and a National Science and Engineering Research Council of Canada (NSERC) Discovery Grant (RGPIN-2025-06669) to A.H. The authors thank the Scialog Fellowship Program from the Research Corporation for Science Advances for facilitating this collaboration.

## AUTHOR CONTRIBUTIONS

A.-K.G. and A.H. conceived the project idea. Q.L. generated the cell line used in the study. T.N. performed the single-molecule experiments and analyzed the data. Z.Q., H.G.-G., and S.V.-H. performed immunofluorescence experiments and analysis. T.N., A.-K.G. and A.H. wrote the manuscript with input from all authors.

