## Supplemental Information for "RhoA effectors ROCK and SLK/LOK oppositely regulate VE-cadherin mobility at endothelial junctions"

### SUPPLEMENTARY FIGURES

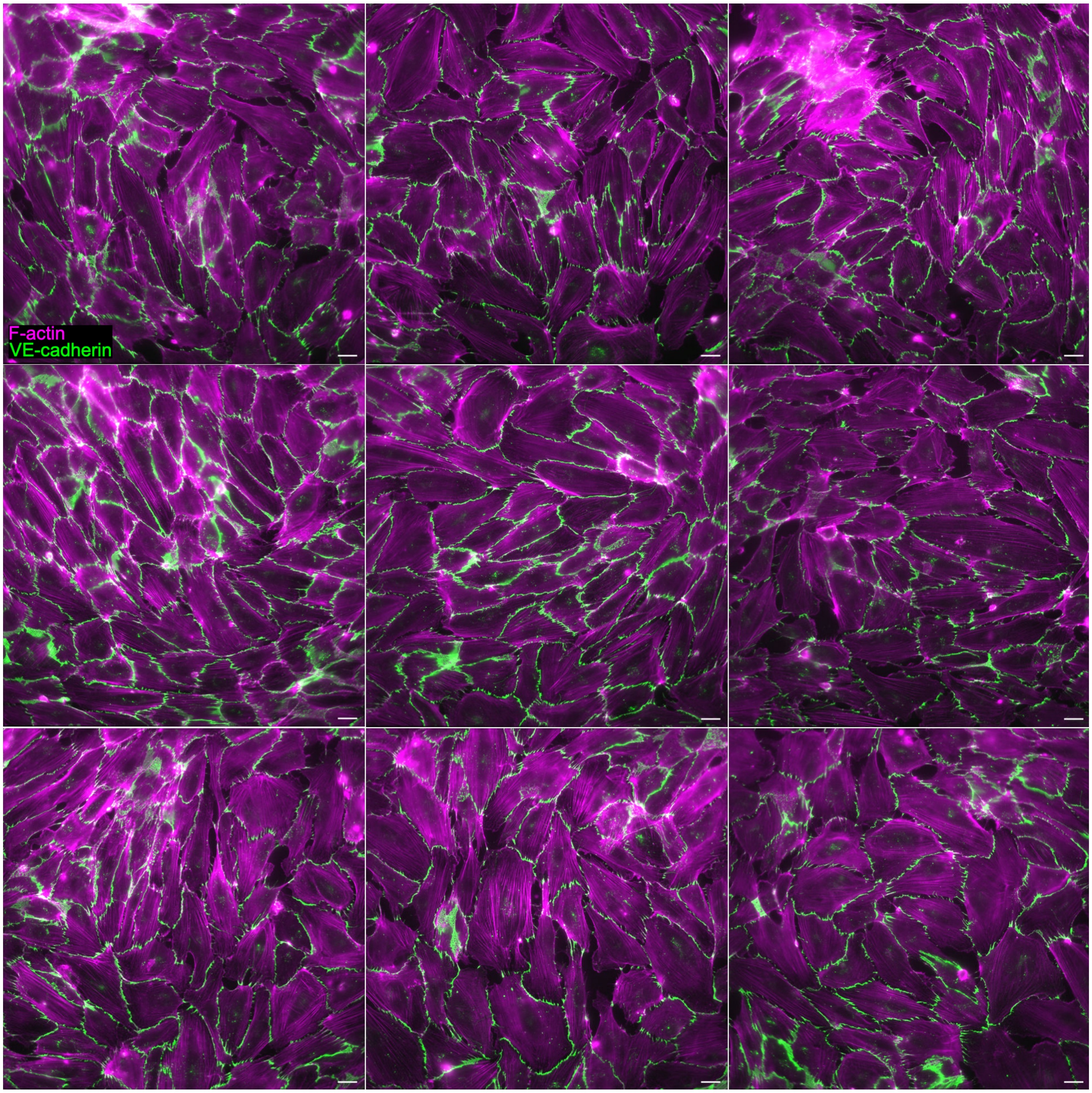


Figure S1. Additional images of untreated HT-HUVEC which were fixed and labeled for F-actin (magenta) and VE-cadherin (green). Scale bars are 20 μm.

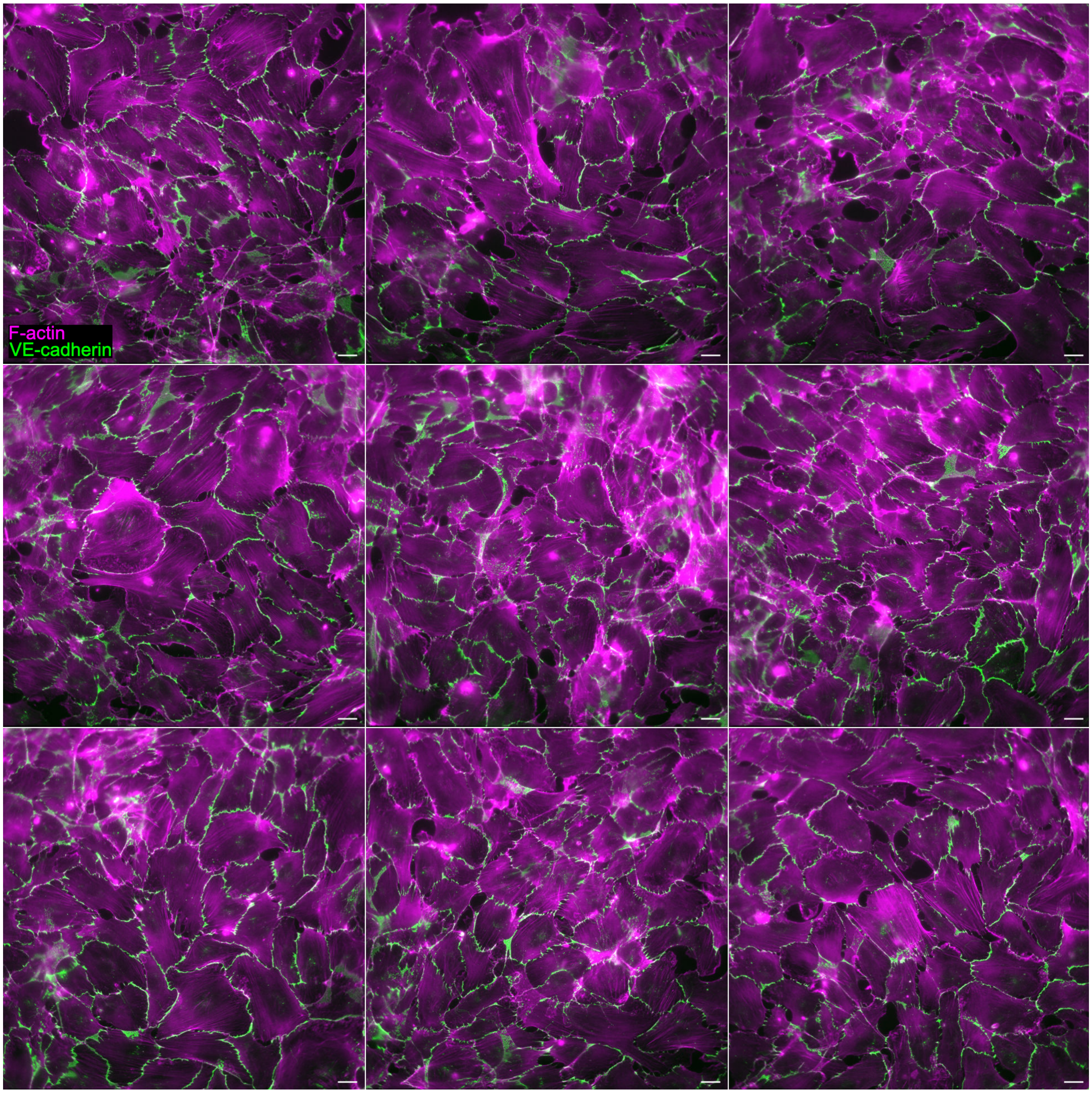


Figure S2. Additional images of HT-HUVEC treated with 20 μM Y-27632 for 2 h, then fixed and labeled for F-actin (magenta) and VE-cadherin (green). Scale bars are 20 μm.

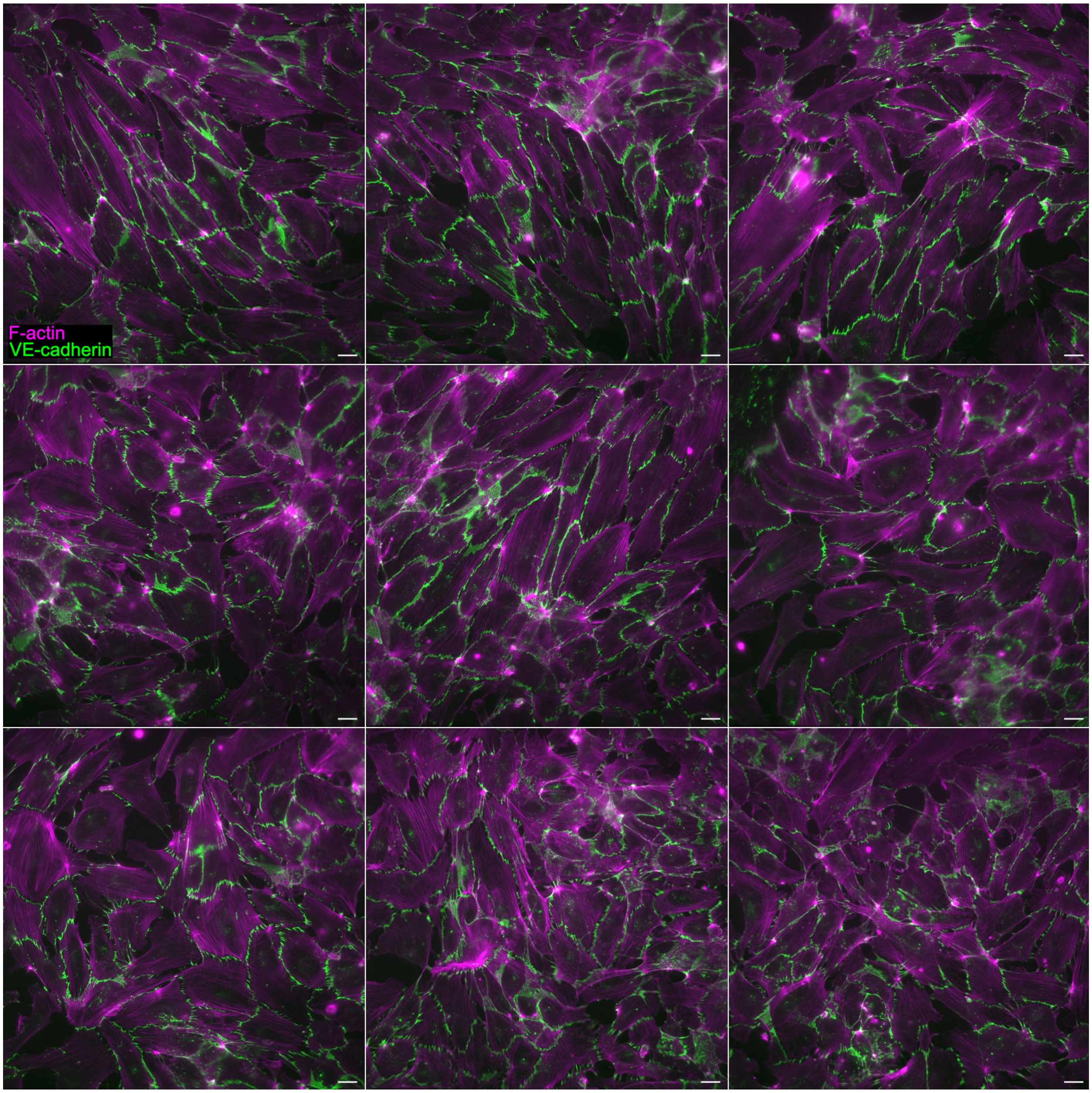


Figure S3. Additional images of HT-HUVEC treated with 5 μM Cpd31 for 2 h, then fixed and labeled for F-actin (magenta) and VE-cadherin (green). Scale bars are 20 μm.
